# A Mechanistic Study on the Cellular Uptake, Intracellular Trafficking, and Antisense Gene Regulation of Bottlebrush Polymer-Conjugated Oligonucleotides

**DOI:** 10.1101/2022.06.15.496353

**Authors:** Lei Zhang, Yuyan Wang, Peiru Chen, Dali Wang, Zheyu Zhang, Ruimeng Wang, Xi Kang, Yang Fang, Hao Lu, Jiansong Cai, Mengqi Ren, Sijia Dong, Ke Zhang

**Author notes:** These authors contributed equally (Lei Zhang, Yuyan Wang).

## Abstract

We have developed a non-cationic transfection vector in the form of a bottlebrush polymer-antisense oligonucleotide (ASO) conjugate. Termed pacDNA (polymer-assisted compaction of DNA), these agents show improved biopharmaceutical characteristics and antisense potency in vivo while suppressing non-antisense side effects. Nonetheless, there still lacks a mechanistic understanding regarding the cellular uptake, subcellular trafficking, and gene knockdown with pacDNA. Here, we show that the pacDNA enters human non-small cell lung cancer cells (NCI-H358) predominantly by scavenger receptor-mediated endocytosis and macropinocytosis, and trafficks via the endolysosomal pathway within the cell. The pacDNA significantly reduces a target gene expression (KRAS) in the protein level but not in the mRNA level, despite that the transfection of free ASOs causes ribonuclease H1 (RNase H)-dependent degradation of KRAS mRNA. In addition, the antisense activity of pacDNA is independent of ASO chemical modification, suggesting that the pacDNA functions as a steric blocker.

**TOC:** 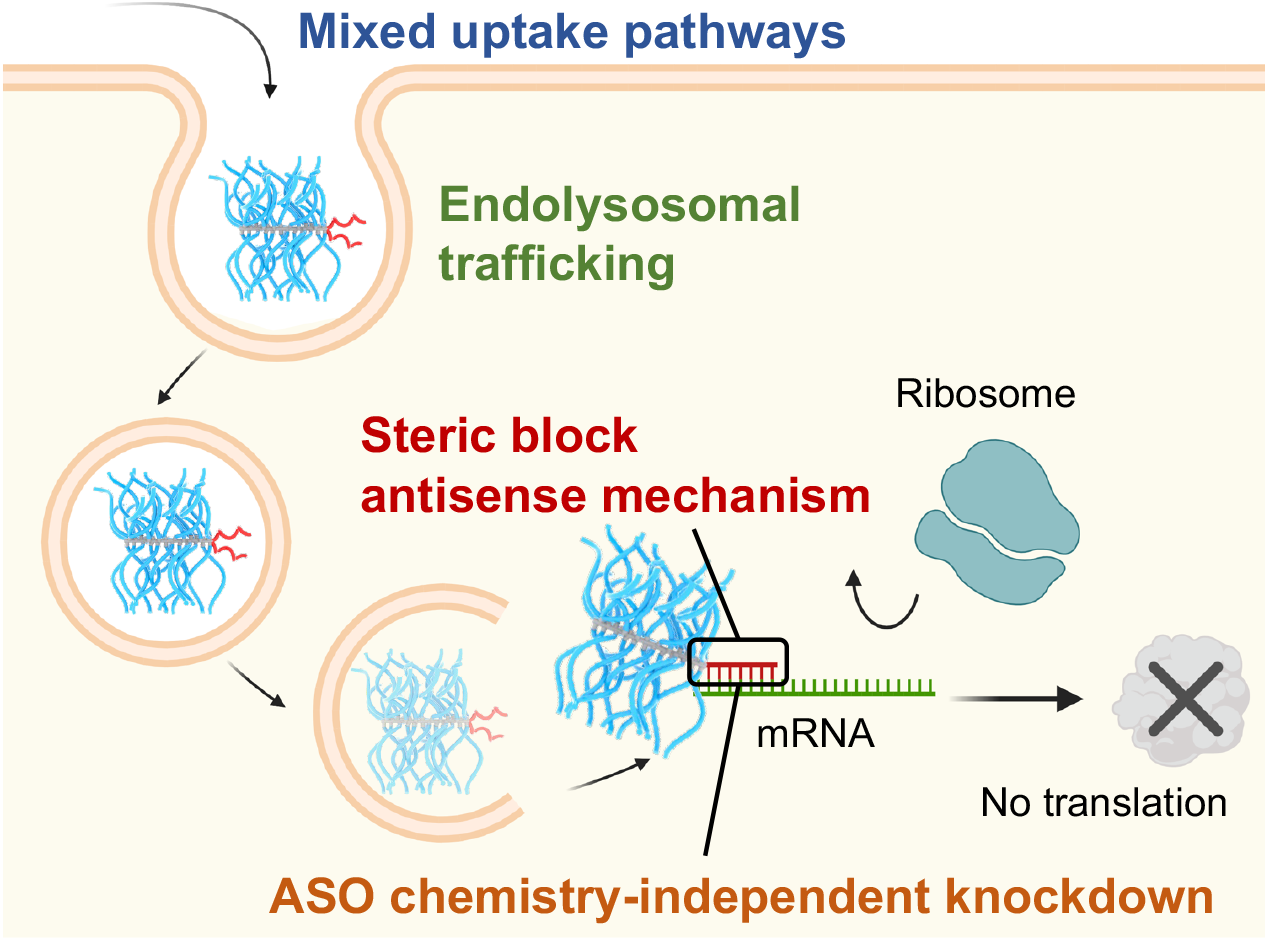

---

Antisense oligonucleotides (ASOs) are widely used in RNA-directed therapies. To date, several ASOs have been approved by the US Food and Drug Administration (FDA) to treat genetic disorders, metabolic diseases, or viral infection, including fomivirsen, mipomersen, nusinersen, eteplirsen, inotersen, golodirsen, and casimersen,^1-3^ with many more in clinical trials. ASOs can modify the expression of the target gene by altering pre-mRNA splicing, degrading mRNA via RNase H, inhibiting translation by steric blocking, and by affecting non-coding RNAs involved in transcriptional and epigenetic regulation.^4^

To guarantee that sufficient ASO molecules reach their cytoplasmic or nuclear targets before being degraded, multiple types of chemical modifications including those of the nucleobase, internucleotide linkage, and the pentose, as well as complete backbone replacements have been developed.^5^ Still, naked ASOs can be cleared rapidly *in vivo*, leading to poor cellular utilization, increased dosage requirement/cost, and narrower therapeutic window. Intracellular delivery systems, which are typically cationic materials (e.g., polymers, peptides, nanoparticles, lipids, ligands, etc.), have been developed to promote cellular uptake and endosomal escape. However, delivery vectors often face a difficult dilemma: features that make cellular transfection more efficient, such as the presence of multiple cationic and hydrophobic groups, often lead to increased toxicity and/or poorer pharmacological properties.^6^ Thus, current efforts in carrier design have often focused on optimizing transfection efficiency within an acceptable toxicity range.

Our group has focused on a different approach to the *in vivo* efficacy problem by prioritizing pharmacological properties of the carrier. The rationale is that, if one can substantially reduce renal clearance, improve plasma pharmacokinetics, and prolong tissue retention, the overall antisense activity *in vivo* can still be greatly enhanced even though the transfection efficiency on the cellular level is moderate. One implication of this philosophy in vector design is that one may do away with polycationic species that drives toxicity, and instead adopt more biologically benign materials that promote blood retention, such as poly(ethylene glycol) (PEG) and zwitterionic polymers.

**Scheme 1.**
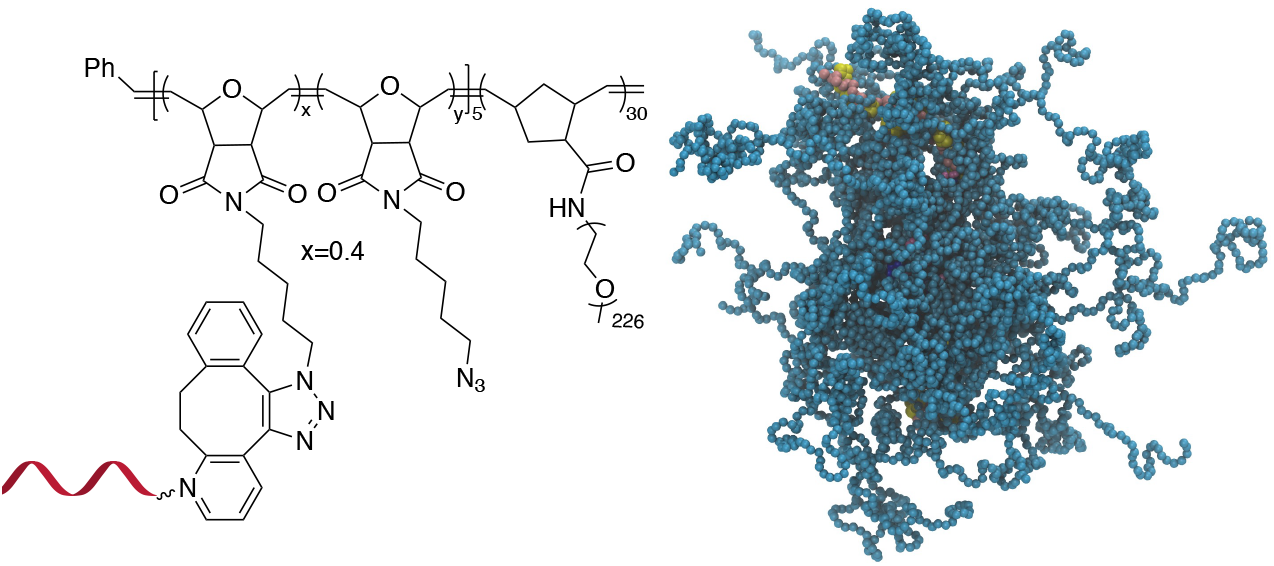
Chemical structure (left) and a coarse-grained molecular dynamics simulation (right) of the pacDNA. In the simulation image, pink/yellow: DNA, cyan: PEG.

Recently, we have developed a novel form of ASO vector, termed pacDNA (polymer-assisted compaction of DNA), which consists of a bottlebrush polymer with a multitude of polyethylene glycol (PEG) side chains and covalently conjugated ASO strands on the polymer backbone (Scheme 1).^7, 8^ The dense arrangement of the PEG side chains endows the ASO with steric selectivity: ASO interactions with proteins are greatly reduced, while hybridization is unaffected in both the kinetic and the thermodynamic sense.^9, 10^ Such selectivity is absent with linear PEG-ASO conjugates and drastically reduces enzymatic degradation and most side effects stemming from specific or non-specific DNA-protein interactions, such as coagulopathy and unwanted innate immune system activation.^11^ Importantly, the pacDNA is significantly more persistent in blood, with 20-50× increase in elimination half-life and 1-2 orders of magnitude increase in area-under-the-curve (AUC_∞_) compared with free DNA. The pharmacological improvements result in substantially enhanced ASO activity *in vivo* in several tumor xenograft models.^12^ In one example where we compared pacDNA with AZD4785, a clinical ASO targeting wild-type KRAS mRNA, pacDNA achieved more pronounced tumor suppression levels than AZD4785 at a fraction (2.5%) of the dosage and with greatly reduced dosing frequency in non-small cell lung carcinoma (NSCLC) mouse models.^13^ In addition, the treatment was free of common deleterious effects such as acute toxicity, inflammation, coagulopathy, and anti-PEG immunity.^13^

When tested *in vitro*, the pacDNA exhibits moderate cellular uptake and can regulate mRNA expression with reasonable efficiency if equipped with an appropriate ASO or siRNA. However, to date, the mechanism for the cellular uptake of the pacDNA and how it regulates protein expression at the molecular level is not well understood. In this study, we explore these aspects using NCI-H358 cells, a human NSCLC cell line harboring the *KRAS*^G12C^ mutation. We anticipate that a mechanistic understanding of the uptake, intracellular trafficking, and gene regulation at the cellular level can not only inform future optimizations of the pacDNA platform but also enable completely novel vector designs based on the *in vivo*-first approach.

The pacDNA used in this study consists on average of ∼30 PEG side chains (10 kDa each) and two ASO strands per molecule, and is synthesized according to prior literature (Scheme S1).^14^ Table S1 provides a summary of the sample nomenclature, ASO sequence, chemical modifications (if any), and assays performed. To study the mechanism for the cellular uptake of pacDNA in NCI-H358 cells, we pre-treated the cells with different endocytosis inhibitors (Table S2) in serum-deprived RPMI-1640 medium for 1 h to block the key pathways of cellular uptake, including lipid raft/caveolae-mediated endocytosis, clathrin-mediated endocytosis, dynamin-mediated endocytosis, and pinocytosis. Next, the pre-treated cells were incubated with Cy3-labeled pacDNA containing phosphodiester (PO) ASO3 (PO pacDNA-3-Cy3) in serum-free medium for 4 h. Flow cytometry analysis revealed depressed fluorescence intensity for cells treated with low temperature (4 °C), dynasore (inhibitor of dynamin^15^), amiloride (inhibitor of epithelial sodium channel, ENaC^16^), and fucoidan (a competitive ligand for scavenger receptor Class A, SR-A^17^), with reductions of ∼97%, ∼38%, ∼45%, and ∼47%, respectively (Figure 2A). The significant reduction in cellular uptake at 4 °C indicates that uptake is predominantly an energy-dependent process as opposed to passive transmembrane diffusion. Sensitivity to amiloride suggests that macropinocytosis plays an important role in the uptake of pacDNA in NCI-H358 cells. Indeed, previous studies report that mutations in Ras proteins can stimulate macropinocytosis in order for cells to use proteins as an amino acid supply.^18^ Cellular uptake is also dependent on dynamin to some extent, which is involved in caveolae- and clathrin-dependent endocytosis. To differentiate the two pathways, cells were treated with filipin III and methyl-β-cyclodextrin (m-β-CD). Both molecules can disrupt the structure of the lipid raft by interfering with cellular cholesterol,^19, 20^ which is required for the formation of caveolae.^21^ The treatment resulted in ∼13% and ∼19% reduction in cell uptake, respectively. On the other hand, treatment with chlorpromazine, which inhibits clathrin-mediated endocytosis by disrupting the assembly and disassembly of clathrin lattices,^22^ reduced the cellular uptake by 21%. SR-A is likely a main receptor for the endocytosis, as treatment with fucoidan blocks roughly half of the uptake, which coincidentally is comparable to the combined contribution of clathrin- and caveolae-dependent endocytosis. SR-A is known to bind to the negatively charged oligonucleotides with its positively charged groove in the collagenous domain.^23^ However, because free DNA is not taken up in high quantities by NCI-H358 cells, it is conceivable that pacDNA improves the adsorption of the DNA onto the plasma membrane, possibly mediated by PEG-cation-membrane coulombic interactions and van der Waals interactions between the hydrophobic polymer backbone and the membrane, leading to more facile SR-A binding to the DNA. Indeed, an anionic form of pacDNA with a negatively charged, hydrophilic backbone undergoes very limited cellular uptake, similar to that of free DNA. Together, these data identify SR-A-mediated endocytosis (both clathrin- and caveolae-dependent pathways) and macropinocytosis as the main mechanisms for the uptake of pacDNA by NCI-H358 cells.

**Figure 1.**
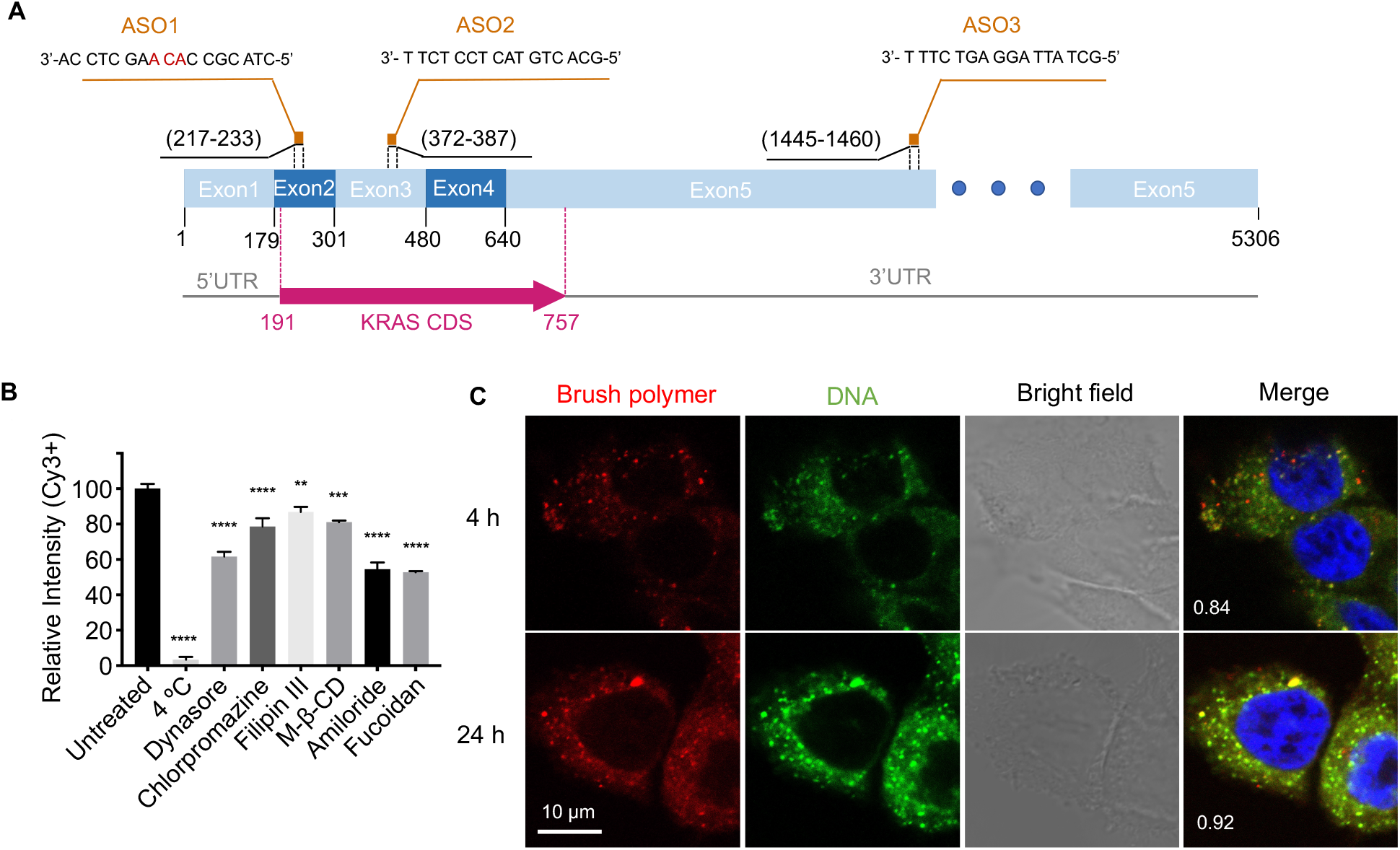
(A) Sequences and binding regions of ASO1, ASO2, and ASO3 to the KRAS transcript isoform b (NM_004985.5). CDS: coding sequence. (B) Uptake of pacDNA by NCI-H358 cells in the presence of various pharmacological blockers. (C) Confocal microscopy of NCI-H358 cells after incubation with dual-labeled pacDNA (red: Cy5-labeled brush polymer; green: Cy3-labeled DNA) for 4 and 24 h. The Manders’ colocalization coefficient is shown in the merged images. The cell nuclei are counter-stained with DAPI and shown in blue.

**Figure 2.**
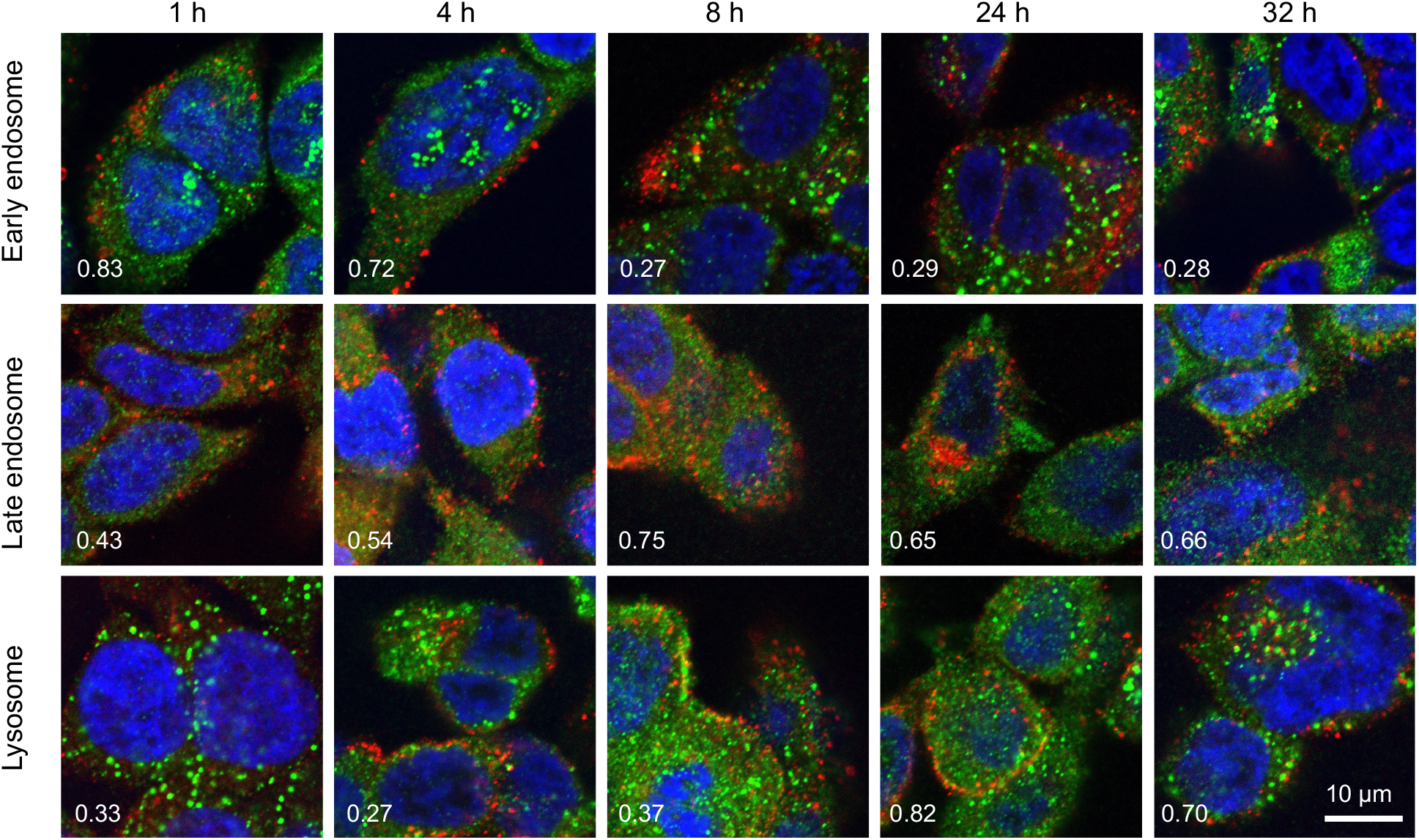
Intracellular trafficking of pacDNA in NCI-H358 cells following different durations of incubation. pacDNA: red; immunofluorescence staining of organelle markers: green. The markers include EEA1 (early endosome), Rab9 (late endosome), and LAMP1 (lysosome). Manders’ colocalization coefficient, shown in the bottom left side of each image, of 0.5 or above indicates substantial colocalization.

Because the pacDNA has two covalently linked components, i.e. the bottlebrush polymer and the oligonucleotide, we first studied whether the two components remain colocalized. To do so, we labeled the bottlebrush polymer with Cy5 and the oligonucleotide with Cy3 (PO Cy5-pacDNA-3-Cy3). The dual-labeled pacDNA was incubated with cells in serum-containing medium, and the co-localizations of the two signals were studied using confocal microscopy. The fluorescence images show punctate patterns in the fluorescent signals, suggesting that the majority of pacDNA remains in endosomal structures. The Cy3 (green) and Cy5 (red) signals are highly colocalized 4 h and 24 h post incubation, with the Manders’ colocalization coefficient of 0.84 and 0.92, respectively (Figure 2C). There are a small number of endosomal patterns where signal colocalization is very poor (red dots), which is likely caused by the movement of those endosomes out of the focal plane between the capture of the Cy3- and Cy5-channel signals. These results suggest that the pacDNA remains largely intact inside the cells for at least 24 h.^7, 14^

The structural stability of the pacDNA in cells indicates that signals from the Cy3 component (DNA) is representative of the intracellular localization of both of the pacDNA components. To study intracellular trafficking following endocytosis, we incubated NCI-H358 cells with Cy3-labeled pacDNA (PO pacDNA-3-Cy3) for different durations of time, followed by immunofluorescence staining to determine their intracellular locations in relation to various protein markers (Figure 3), including early endosome antigen 1 (EEA1, early endosome),^24^ Ras-related protein 9 (Rab9, late endosome),^25^ and lysosomal-associated membrane protein 1 (LAMP1, lysosome).^26^ After 1 h of incubation, pacDNA predominantly colocalizes with early endosome (Manders’ colocalization coefficient of 0.83). This value decreases to 0.72 at 4 h and to 0.29 at 8 h post-incubation, suggesting that the pacDNA is trafficked away from the early endosome. In the meantime, colocalization with late endosome increases from 0.43 (1 h) to 0.75 (8 h). Colocalization of pacDNA and LAMP1 also increases from 0.33 to 0.82 in 24 h, suggesting a significant fraction of pacDNAs are transported to the lysosomes. We performed the same set of experiments in serum-deprived conditions, and pacDNA exhibited a similar trafficking pathway (Figure S1). Collectively, pacDNA primarily adopts the conventional endolysosomal route of trafficking after entering NCI-H358 cells, irrespective of the presence of serum proteins in the culture media.

**Figure 3.**
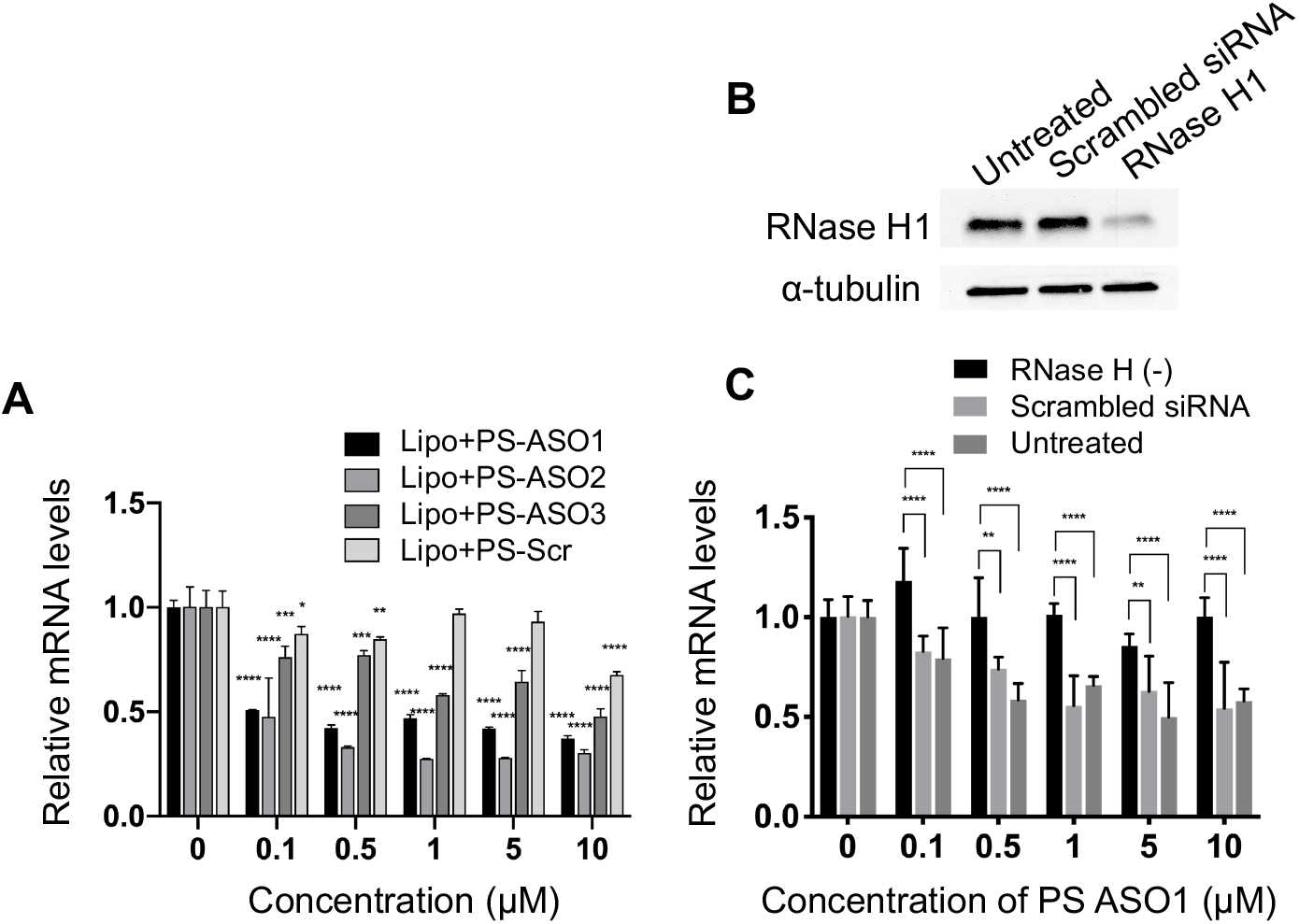
(**A**) Lipofectamine2k-assisted transfection of PS ASOs for 24 h causes significant sequence-dependent reduction in KRAS mRNA levels in NCI-H358 cells. (**B**) Western blot of NCI-H358 cell lysates after cells were treated with siRNA to knockdown RNase H1. (**C**) Reduction of RNaseH1 eliminates the gene knockdown effect of PS ASO1, suggesting that free PS ASO1 depends on RNase H1 for target mRNA degradation. qPCR results are shown in mean ± standard deviation from at least three individual experiments. All results are normalized to β-actin mRNA. \**P*<0.05, \*\**P*<0.01, \*\*\**P*<0.001, and \*\*\**P*<0.0001 (one-way ANOVA, Tukey’s test).

To probe how the pacDNA inhibits the expression of target protein, we selected three antisense ASOs that have been shown effective in lowering KRAS mRNA. These ASOs target the G12C mutation site in exon 2 (ASO1),^27^ a bulge in exon 3 (ASO2),^28^ and a segment in the 3’ UTR of KRAS mRNA (ASO3, Figure 1A and Table S2).^29^ We first confirmed the antisense activity of the free ASOs with full phosphorothioate (PS) modifications. Lipofectamine2k, a cationic liposomal transfection agent, was used to form lipoplexes with the ASOs, which were incubated with NCI-H358 cells for 24 h before mRNA levels were determined by quantitative real-time polymerase chain reaction (qRT-PCR) (Figure 3A). All three ASOs showed apparent reductions in the KRAS transcript levels in the tested concentration range (0.1-10 μM), while a scrambled ASO did not result in apparent knockdown. The antisense activity was in the order: ASO2 > ASO1 > ASO3. The downregulation of KRAS mRNA is likely mediated by RNase H1, an endonuclease that degrades the RNA of an RNA/DNA heterocomplex.^30^ When RNase H1 was depleted by an siRNA for 3 days in NCI-H358 cells (Figure 3B), subsequent treatment with lipofectamine2k-complexed PS-ASO1 did not result in KRAS mRNA downregulation(Figure 3C).

Next, pacDNA formulations of the ASOs 1-3 (named correspondingly as pacDNA 1-3, both PO and PS versions) were prepared and tested. We first screened the pacDNAs for inhibition of cellular growth (Figure S2). The least effective sample, pacDNA-2, was removed from subsequent KRAS knockdown studies. Next, NCI-H358 cells were treated with the PO form of pacDNA-1 and pacDNA-3 for 24 h and 48 h, but no significant reduction of KRAS mRNA was detected at either time points by qRT-PCR (Figure 4A). Expectedly, knocking down RNase H1 beforehand had no effect on KRAS mRNA levels in response to pacDNA treatment (Figure 4B). These results suggest that the pacDNA does not significantly affect the mRNA level of the target gene, despite the ASO being RNase H-acceptable. However, KRAS protein levels were notably reduced in a dose-dependent manner when cells were treated with both PS and PO forms of pacDNAs for 72 h (Figure 5), while scrambled pacDNA controls (pacDNA-scr in both PO and PS forms) did not lead to KRAS downregulation. Knockdown efficiency was between 17-40% at 1 μM ASO concentration as determined by densitometry analysis of western blots. Interestingly, although free PS ASO1 was able to significantly reduce KRAS mRNA transcript levels, its ability to affect KRAS protein levels is comparable to PS pacDNA-1. These results suggest that pacDNA in general inhibits protein expression via the steric block mechanism, for example by disrupting the regulatory functions of 3’ and 5’ UTR, by preventing ribosome assembly, or by steric blockade of the start codon/coding sequence. The observation that pacDNAs with PO or PS ASOs do not exhibit RNase H-dependent mRNA cleavage may be attributed to the bottlebrush component, which blocks RNase H from accessing the mRNA/ASO heteroduplex. One implication of this mechanism is that the gene knockdown for pacDNA becomes independent of ASO chemistry. To test this hypothesis, we furnished the pacDNA with a fully 2’-*O*-methyl-modified ASO1 (OMe pacDNA-1). This modification is known to be less toxic than PS and have higher affinity for their target sequence, but is more RNA-like and cannot induce RNase H-dependent degradation. Indeed, Lipofectamine2k-assisted transfection using PS OMe ASO1 did not reduce the KRAS mRNA levels (Figure S5), neither did OMe pacDNA-1 (Figure 6A-C). With western blotting, however, significant reduction (41%) of KRAS protein expression was observed in cells treated with OMe pacDNA-1 at 1 μM ASO for 72 h (Figure 6D). The reduction in KRAS protein levels associated with pacDNA treatment inhibits cellular growth (Figure 6E). Given that pacDNAs reduce target proteins levels but does not lower target mRNA levels, and that the effect is ASO chemistry-independent, it is likely that the pacDNA works as a steric blocker in the translation process.

**Figure 4.**
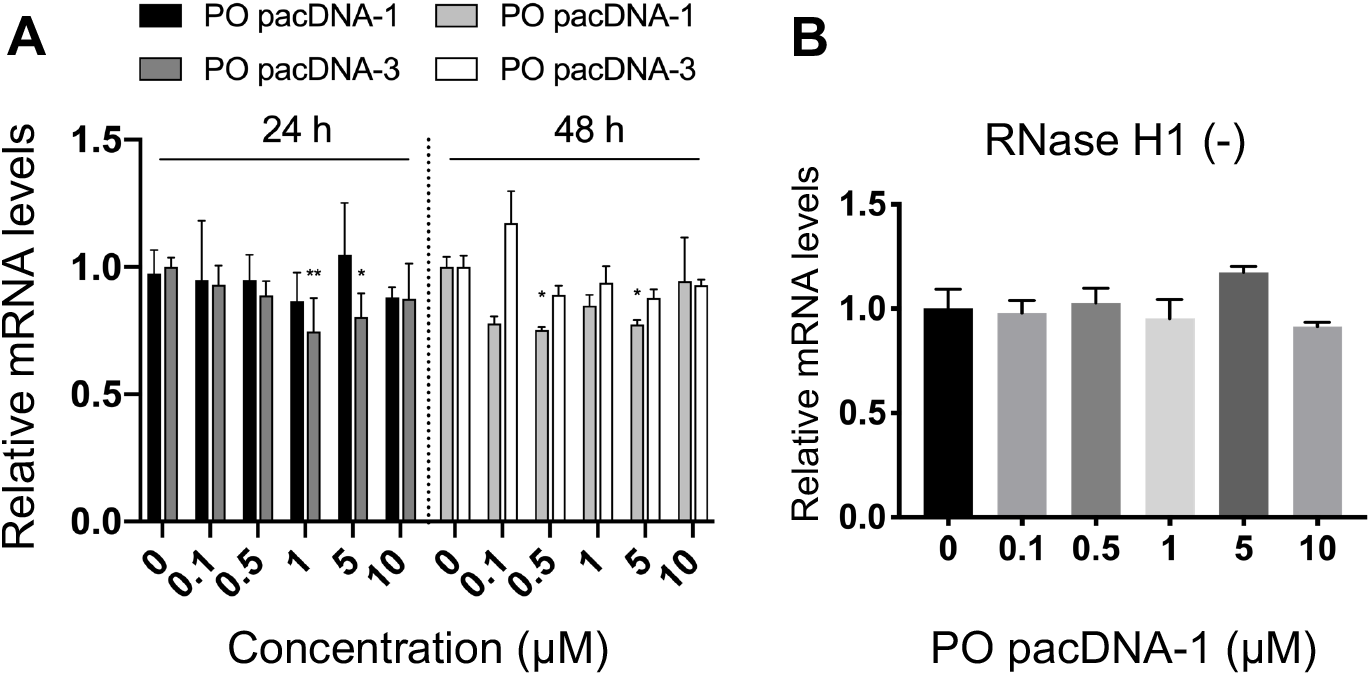
(**A**) Relative KRAS mRNA levels upon treatment with pacDNA containing different ASOs for 24 h or 48 h, showing no significant down-regulation. (**B**) Pre-knockdown of RNaseH1 has no effect on KRAS mRNA levels following incubation with PO pacDNA-1 for 24 h. The qRT-PCR results are shown in mean ± standard deviation from at least three individual experiments. All results are normalized to β-actin mRNA. \**P*<0.05, \*\**P*<0.01 (one-way ANOVA, Tukey’s test).

**Figure 5.**
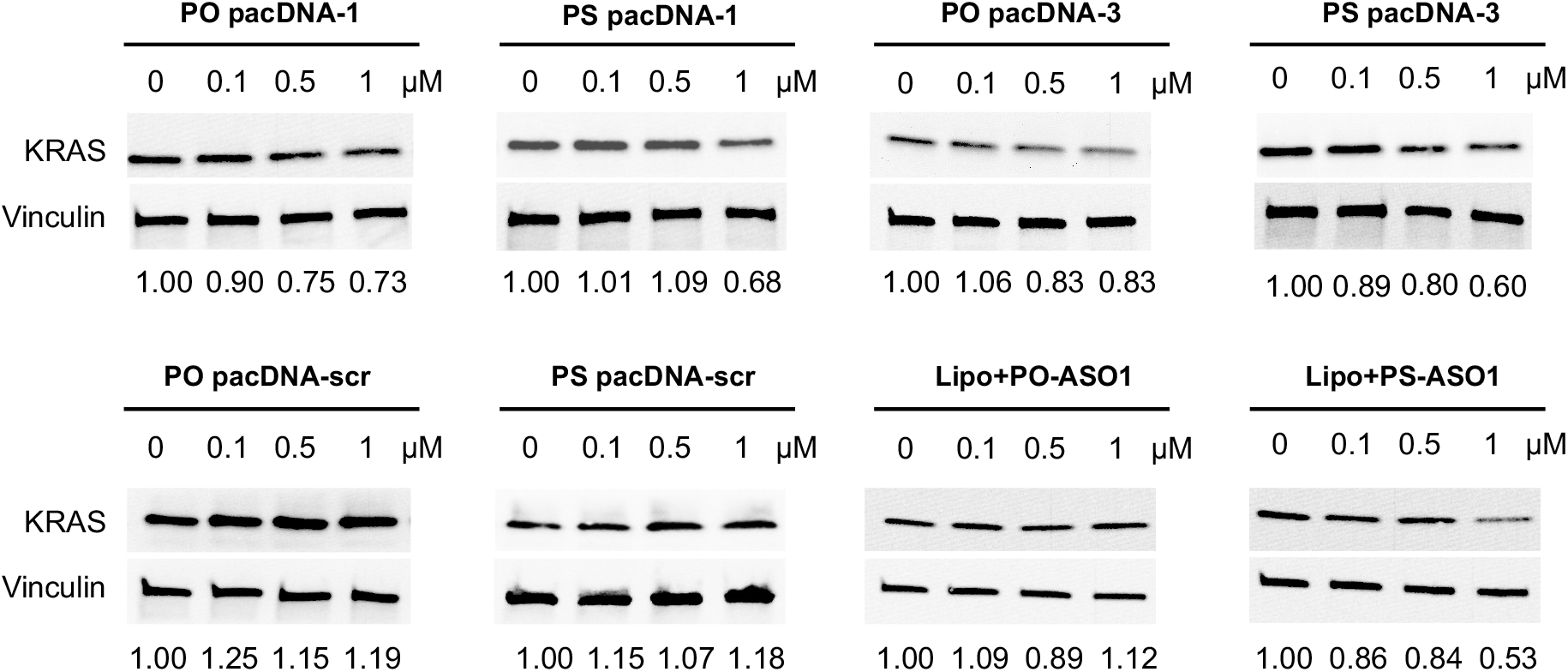
Western blot of NCI-H358 cell lysates after cells were incubated with pacDNA samples and controls at a concentration range of 0-1.0 μM for 72 h. The relative KRAS protein expression levels are shown below the blot images after normalization to vinculin.

**Figure 6.**
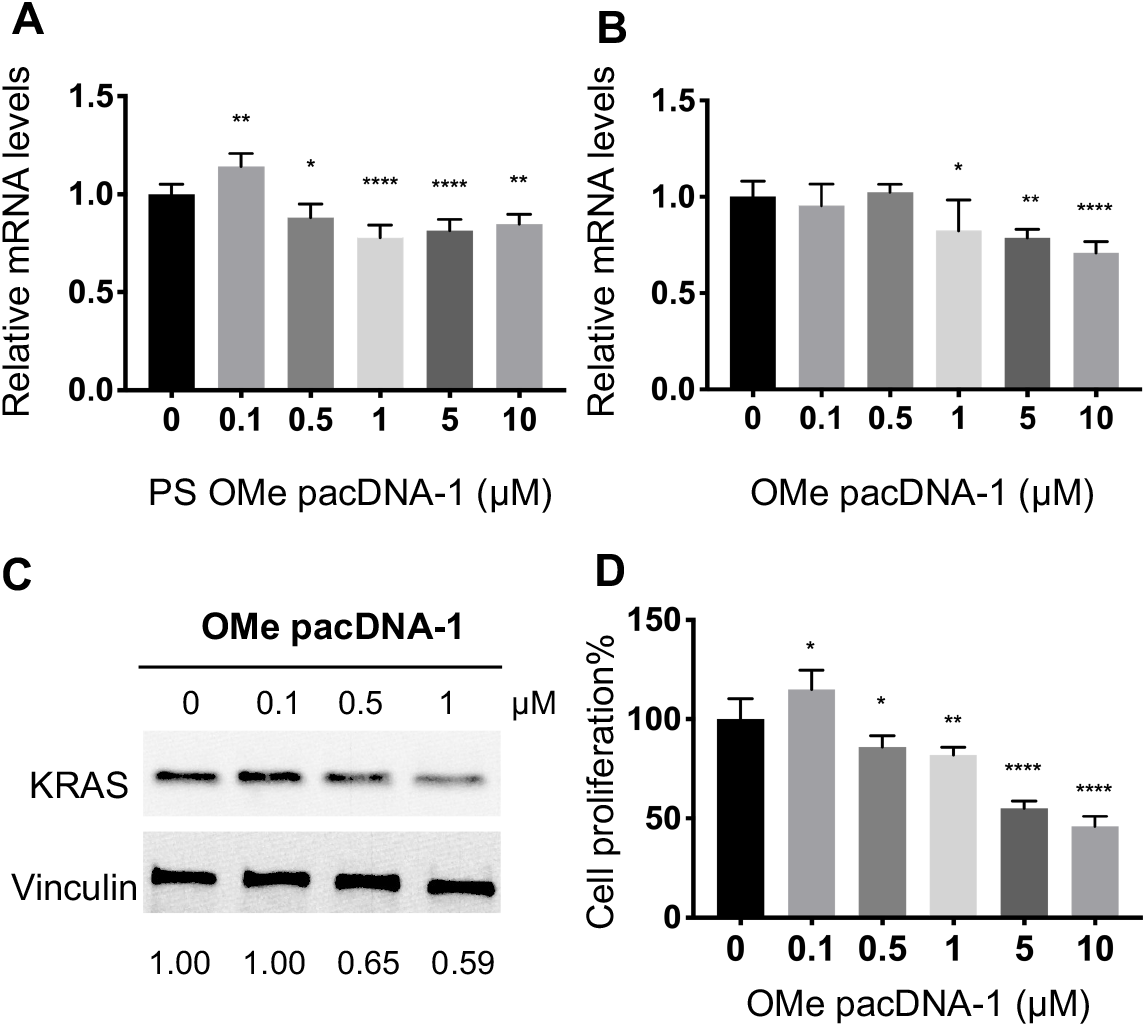
(**A-B**) Both Lipofectamine2k-assisted transfection of PS OMe ASO1 and unformulated OMe pacDNA-1 do not reduce KRAS mRNA levels. The qRT-PCR results are shown in mean ± standard deviation from at least three individual experiments. All results are normalized to β-actin mRNA. \**P*<0.05, \*\**P*<0.01 (one-way ANOVA, Tukey’s test). (**C**) Western blot analysis of NCI-H358 cells after incubation with OMe pacDNA-1 at concentrations 0-1 μM for 72 h. The expression levels of KRAS normalized to vinculin are shown below the blot images. (**D**) Viability of NCI-H358 cells treated with OMe pacDNA-1 for 96 h as determined by an MTT assay. Error bars denote the standard deviation of five individual experiments.

## Conclusions

In summary, we demonstrate that the pacDNA construct enters NCI-H358 cells predominantly via SR-A-mediated endocytosis and macropinocytosis. After endolysosomal trafficking, a fraction of the pacDNA gains access to the cytosol, where it causes translational arrest of the target mRNA by steric blocking. The activity of the pacDNA is not dependent on the ASO chemistry; non-RNase H active ASO modifications can still cause target gene downregulation as long as they retain target binding affinity. We anticipate these fundamental understanding of how the pacDNA functions in vitro will provide the foundation for more in-depth mechanistic explorations and disease-specific structural optimizations of the pacDNA.

## Author Contributions

K.Z. and L.Z. devised the experiments and wrote the manuscript. L.Z. and Y.W. conducted the synthesis of materials, purification, and material/biological characterizations. All other authors contributed to material synthesis, purification, molecular dynamics simulation, and/or discussion of the results. All authors edited the manuscript.

## Acknowledgements

This work was supported by the National Institute of General Medical Sciences (1R01GM121612), the National Cancer Institute (1R01CA251730), and the National Science Foundation (DMR award number 2004947).

## Supplementary Information

### Materials

ω-Amine PEG methyl ether (Mn=10 kDa, PDI=1.05) was purchased from JenKem Technology, USA. Dibenzocyclooctyne-N-hydroxysuccinimidyl ester (DBCO-NHS) was purchased from Sigma-Aldrich Co., USA. Phosphoramidites and supplies for DNA synthesis were obtained from Glen Research Co., USA. NCI-H358 human non-small cell lung cancer cell line was purchased from American Type Culture Collection (Rockville, MD, USA). All other materials were obtained from Fisher Scientific Inc., USA, or VWR International LLC., USA, and used as received unless otherwise indicated.

### Instrumentation

*N,N*-dimethylformamide (DMF) gel permeation chromatography (GPC) was performed on a TOSOH EcoSEC HLC-8320 GPC system (Tokyo, Japan) equipped with a TSKGel GMHHR-H, 7.8×300 mm column and RI/UV-Vis detectors. HPLC-grade DMF with0.05 M LiBr was used as the mobile phase, and samples were run at a flow rate of 0.4 mL/min. GPC calibration was based on polystyrene standards (706 kDa, 96.4 kDa, 5970 Da, 500 Da). Aqueous GPC measurements were carried out on a Waters Breeze 2 GPC system equipped with an Ultrahydrogel™ 1000, 7.8×300 mm column and three Ultrahydrogel™ 250, 7.8×300 mm columns and a 2998 PDA detector for the separation of as-synthesized polymers from monomers/oligonucleotides. Sodium nitrate solution (0.1 M) was used as the eluent running at a flow rate of 0.8 mL/min. To purify the oligonucleotides, reversed-phase HPLC was performed on a Waters (Waters Co., MA, USA) Breeze 2 HPLC system coupled to a Symmetry^®^ C18 3.5 μm, 4.6×75 mm reversed-phase column and a 2998 PDA detector, using TEAA buffer (0.1 M) and HPLC-grade acetonitrile as mobile phases. MALDI-TOF MS measurements were performed on a Bruker Microflex LT mass spectrometer (Bruker Daltonics Inc., MA, USA).

### Oligonucleotide synthesis

Oligonucleotides including modifications were synthesized on a Model 391 DNA synthesizer (Applied Biosystems, Inc., Foster City, CA) using standard solid-phase phosphoramidite methodology. DNA were cleaved from the CPG support using aqueous ammonium hydroxide (30% NH_3_ basis) at room temperature for 18 h. OMe-modified strands were synthesized on Universal Support III PS CPG, cleaved by treating with 2 M ammonia in methanol at room temperature for 60 minutes, and deprotected using aqueous ammonium hydroxide (30% NH_3_ basis) at room temperature for 18 h. All strands were purified by reversed-phase HPLC. The successful synthesis of all sequences was confirmed by MALDI-TOF MS.

### Synthesis of bottlebrush polymer

The diblock bottlebrush polymer (**3**, Scheme S1) was synthesized via ring-opening polymerization of norbornenyl bromide (**1**) and norbornenyl PEG (**2**), following by azide substitution. The synthesis of norbornenyl bromide, norbornenyl PEG, and modified 2^nd^ generation Grubbs’ catalyst has been described in our previous publications.^1-2^ Typically, a solution of norbornenyl bromide (**1**, 5 equiv.) in deoxygenated dichloromethane was added into a Schlenk flask under N_2_. The solution was cooled to -20 °C in an ice-salt bath, to which modified Grubbs’ catalyst (1 equiv.) in deoxygenated dichloromethane was added. The reaction mixture was stirred vigorously for 30 min until thin-layer chromatography (TLC) confirmed the complete consumption of the monomer. Then, a solution of **2** (30 equiv.) in deoxygenated dichloromethane was added to the reaction. The reaction mixture was further stirred for 6 h, before addition of several drops of ethyl vinyl ether (EVE) to remove the chain-end catalyst. Note that all solution was added by gastight syringe to reduce the amount of oxygen introduced to the reaction. The mixture was stirred overnight, and the product was precipitated into cold diethyl ether 3× and dried under vacuum. The as-synthesized polymer was treated with an excess of sodium azide in DMF overnight at room temperature to give the azide-functionalized bottlebrush polymer, **3**. The product was dialyzed against Nanopure^™^ water for 24 h, lyophilized, re-dissolved in Nanopure^™^ water, and injected into an aqueous GPC for collection of the fractions containing the bottlebrush polymer. The final polymer was desalted using a NAP-10 column (G.E. Healthcare, IL, USA). DMF-GPC analysis determines the bottlebrush polymer with Mn = 290 kDa, Mw = 390 kDa, PDI = 1.34.

### Synthesis of pacDNA

In a typical procedure, azide-functionalized bottlebrush polymer **3** (50 nmol) was dissolved in 100 μL Nanopure^™^ water, to which DBCO-modified DNA (or chemically modified forms) was added (2.2 equiv. to N_3_, 50 μL aqueous solution). The reaction mixtures were shaken gently for 24 h at 50 °C on an Eppendorf Thermomixer. Thereafter, aqueous GPC was used to isolate the conjugation product from unreacted DNA. The conjugates were desalted using a NAP-10 column and lyophilized to yield a white powder (or red powders for Cy3-labeled pacDNA).

### Synthesis of dual-labeled pacDNA

To simultaneously label the bottlebrush polymer and DNA strands for *in vitro* fluorescence tracking, the azide-functionalized polymer was first labelled with Cy5 before coupling to Cy3-labeled DNA. Specifically, polymer **3** (50 nmol) and alkyne-modified Cy5 (150 nmol, 150 μL of 1 mM stock solution in DMSO) were mixed in Nanopure^™^ water (2 mL), followed by the addition of the catalyst system (CuSO_4_·5H_2_O, 40 nmol; tris(3-hydroxypropyltriazolylmethyl)amine known as THPTA, 50 nmol; sodium ascorbate, 250 nmol). After 12 hours of stirring, the reaction mixture was dialyzed against Nanopure^™^ water to remove small molecular residuals and lyophilized. The Cy5-labeled bottlebrush polymer was then coupled to Cy3-conjugated DNA under the same conditions as described above, purified by aqueous GPC, desalted, and lyophilized to yield a purple powder.

### Cell culture

NCI-H358 cells were cultured in RPMI-1640 medium supplied with 10% fetal bovine serum (FBS, Gibco, USA), 100 units/mL of penicillin and 100 µg/mL of streptomycin (Gibco, USA) at 37 °C in a humidified atmosphere containing 5% CO_2_.

### Cellular uptake

Cellular uptake kinetics was evaluated using flow cytometry. Cells were seeded in 24-well plates at 3.0×10^5^ cells per well in 1 mL complete RPMI-1640 medium and cultured for 24 h at 37 °C with 5% CO_2_. pacDNA-3-Cy3 (2 μM equiv. of DNA) was dissolved in RPMI culture medium containing 10% FBS, and cells were further incubated at 37 °C for 10 min through 48 h. In a parallel study, cells were incubated with pacDNA in RPMI culture medium without FBS. At predetermined time points, cells were washed with PBS before being treated with trypsin-EDTA (Gibco, USA). Thereafter, 2 mL of PBS was added to each culture well, and the solutions were centrifugated for 5 min (1000 rpm). Cells were then resuspended in 0.5 mL of PBS for flow cytometry analysis on an Attune NxT flow cytometer (Thermo Fisher Scientific, USA). The fluorescence signals were collected using 455 nm as the excitation laser, with the BL2 channel (574/26 nm) as the emission filter. For each measurement, 3.0×10^4^ gated events were collected. The experiments were carried out in triplicates.

For confocal microscopy, cells were seeded on a glass slide of 12 mm in diameter (Marienfeld Superior) in a 24-well plate (Fisher Scientific, USA) at 3×10^5^ cells per well and cultured for 24 h. The dual-labeled Cy5-pacDNA-3-Cy3 mixed with 0.5 mL full medium (2 μM equiv. of DNA) was added, and the cells were further incubated for 4 h or 24 h. Cells were then washed with PBS, fixed with 4% paraformaldehyde (PFA), and counter-stained with Hoechst 33342 for 10 min. Cells were washed with PBS 3×, mounted to the glass slides with Fluoromount-G™ mounting medium (Thermo Fisher Scientific, USA), and dried overnight. The samples were then imaged on an LSM-700 confocal laser scanning microscope (Carl Zeiss Ltd., Cambridge, UK). Images were taken at excitation wavelengths of 350 nm (Hoechst 33342), 532 nm (Alexa Fluor 488), and 640 nm (Alexa Fluor 488). To quantify the colocalization of the two fluorescence signals of pacDNA, Image J (Fiji contributors) with a Coloc2 plugin was used to calculate the Manders colocalization coefficient according to the protocols provided by the software authors, where G represents green pixel signals (pseudocolor for Cy3) and R represents red pixel signals (Cy5). The threshold value was kept identical for all images analysed and the M1 coefficient was used for comparisons.

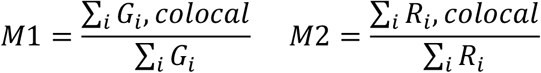

### Pharmacological inhibition

Cells were seeded in 24-well plates at 3×10^5^ cells per well overnight before being pre-treated with 300 μL of RPMI-1640 medium that contains different concentrations of chemical blockers for 1 h (Table S2). These blockers include fucoidan (50 μg/mL, Sigma Aldrich Fine Chemicals Biosciences), filipin III (2.5 μg/mL, Cayman Chemical), chlorpromazine (5 μg/mL, Tokyo Chemical Industry), amiloride (500 μg/mL, Cayman Chemical), dynasore (120 μM, MilliporeSigma™), and methyl-β-cyclodextrin (M-β-CD) (10 mM, Sigma-Aldrich). After removing the inhibitor-containing medium, 0.5 mL of serum-containing medium or sole RMPI-1640 medium that contains the same inhibitor at the original concentration and 2 μM of pacDNA-3-Cy3 were subsequently added to the cells. After 4 h of incubation, the medium was removed, and the cells were rinsed with PBS and trypsinized. Cells were then resuspended in 0.5 mL of PBS for flow cytometry analysis on an Attune NxT flow cytometer (Thermo Fisher Scientific, USA). The Cy3 fluorescence signals were collected using blue laser (455 nm) as the excitation light with BL2 channel (574/26 nm) as the emission filter. For each measurement, 3.0×10^4^ gated events were collected. The experiments were carried out in triplicates.

### Immunofluorescence staining of cells

Cells were seeded on a glass slide of 12 mm in diameter (Marienfeld Superior) in a 24-well plate (Fisher Scientific, USA) at 3×10^5^ cells per well overnight before being incubated with 0.5 mL FBS-containing medium containing 2 μM of pacDNA-3-Cy3 for 1 h through 32 h. In a parallel study, 2 μM of pacDNA-3-Cy3 in RPMI-1640 medium without FBS was used. After treatment, cells were washed, fixed with 4% paraformaldehyde (PFA), and permeated with 1% Triton X-100 (Sigma-Aldrich) for 10 min. Cells were blocked with 1% BSA in PBS for 1 h at room temperature, before being stained with primary antibodies (in PBS with 1% BSA) overnight at 4 °C. The primary antibodies include those antibodies against EEA1 (Invitrogen, Cat. No. MA5-14794) (1:500), Rab9 (Invitrogen, Cat. No. MA3-067) (1:500), and Lamp1 (Abcam, Cat. No. ab24170) (1:1000). After rinsing with 0.05% Tween-20 in PBS, cells were stained with Alexa Fluor 488-conjugated goat anti-rabbit IgG (H+L) secondary antibody (Invitrogen, Cat. No. A11008) (for EEA1 and Lamp1), or Alexa Fluor 488-conjugated goat anti-mouse IgG secondary antibody (Invitrogen, Cat. No. A11029) at 1 μg/mL (1% BSA in PBS) for 1 h at room temperature. Cells were counter-stained with Hoechst 33342 for 10 min, washed with PBS, mounted to the glass slides with Fluoromount-G™ mounting medium (Thermo Fisher Scientific, USA), and dried overnight. The samples were then imaged on an LSM-700 confocal laser scanning microscope (Carl Zeiss Ltd., Cambridge, UK). Images were taken at excitation wavelengths of 350 nm (Hoechst 33342) and 488 nm (Alexa Fluor 488). To quantify the colocalization between fluorescence signals of pacDNA and cellular compartments, Image J (Fiji contributors) with a Coloc2 plugin was used to calculate the Manders colocalization coefficient according to the protocols provided by the authors, where R represents red pixel signals (Cy3) and G represents green pixel signals (Alexa Fluor 488). Manders’ tM2 (above autothreshold of red channel) coefficient was use to indicate the degree of co-localization.

### MTT assay

The cell proliferation inhibition effect of pacDNA was evaluated with the MTT assay against NCI-H358 cells. Briefly, NCI-H358 cells were seeded into 96-well plates at 2.5×10^4^ cells per well in 200 μL full medium and cultured for 24 h. The cells were then treated with pacDNA at varying concentrations of DNA (0.1 through 10 μM). Cells treated with vehicle (PBS) were used as a negative control. After 96 h of incubation, 20 μL of 5 mg/mL MTT stock solution in PBS was added to each well. The cells were incubated for another 4 h, and the medium containing unreacted MTT was removed carefully. The resulting blue formazan crystals were dissolved in 200 μL DMSO per well, and the absorbances (490 nm) were measured on a BioTek® SynergyTM Neo2 Multi-Mode microplate reader (BioTek Inc., VT, USA).

### Quantification of mRNA levels by qRT-PCR

NCI-H358 cells were plated at a density of 3.0×10^5^ cells per well in 24-well plates and cultured overnight. Cells were incubated with pacDNA (0.1 through 10 μM), or Lipofectamine2k-complexed DNA (0.1 through 10 μM) in full medium for specified durations of time. Lipofectamine2k was first mixed with PS ASOs in serum-free medium for 15 min before addition to full medium at a concentration of 1 μg/mL. A two-step qRT-PCR method was used to quantify KRAS mRNA levels. After incubation, the total RNA was extracted using the Trizol reagent (Invitrogen, Thermo Fisher Scientific) following manufacture-suggested protocols. The RNA concentration was determined using a NanoDrop 2000 spectrophotometer (Thermo Scientific). Total RNA (300 ng) was reverse-transcribed to cDNA using the SuperRT cDNA Synthesis kit (CWBIO) at 42 °C for 30 min and 85 °C for 5 min. The cDNA was amplified with SsoAdvanced Universal SYBR Green Super Mix on a Bio-Rad CFX96 Touch System. The results were normalized to β-actin expression. The primer sequences used were: KRAS forward-GAC ACA AAA CAG GCT CAG GAC TT, reverse-TCT TGT CTT TGC TGA TGT TTC AAT AA; β-actin forward-CGG ACT ATG ACT TAG TTG CGT TAC A, reverse-GCC ATG CCA ATC TCA TCT TGT. A second primer set was use to verify the result by normalization to GAPDH expression: KRAS forward-GCC TGC TGA AAA TGA CTG AAT ATA, reverse-TTA GCT GTA TCG TCA AGG CAC TC, GAPDH forward-AAT CCC ATC ACC ATC TTC CA, reverse-TGG ACT CCA CGA CGT ACT CA. All qRT-PCR experiments were performed at least in triplicates and the results were averaged. Fold difference in relative changes in gene expression is according to the 2^−ΔΔCt^ formula as described by Livak et al.^3^

### Western blotting

Cells were seeded at a density of 2.0×10^5^ cells in a 24-well plate overnight, and incubated with pacDNA or Lipofectamine2k-complexed DNA in serum-containing medium for 72 h. Thereafter, whole-cell lysates were collected in 100 μL of RPIA lysis and extraction buffer (Thermo Scientific, USA) containing Halt™ protease and phosphatase inhibitor cocktail (Thermo Scientific, USA) following manufacturer’s protocols. Protein content in the extracts was quantified using a bicinchoninic acid protein assay kit (Thermo Fisher Scientific, MA, USA). Equal amounts of proteins (20 μg per lane) were separated on 4-20% gradient SDS–polyacrylamide gel electrophoresis and electro-transferred to a nitrocellulose membrane. The membranes were then blocked with 3% bovine serum albumin in tris-buffered saline supplemented with 0.05% Tween 20 (TBST) and further incubated with primary antibodies at 4°C overnight, including KRAS antibody (Cat. No. NBP2-45536, Novus Biologicals, CO, USA) (1:1000 dilution), phospho-p44/42 MAPK (Erk1/2) (Thr202/Tyr204) antibody (Cell Signalling Technology, MA, USA) (1:1000 dilution) primary, and vinculin monoclonal antibody, clone hVIN-1 (Cat. No. V9131, Sigma Aldrich, USA) (1:200 dilution). After washing with TBST 3×, membranes were further incubated with anti-mouse IgG, HRP-linked secondary antibody (Cell Signalling Technology, MA, USA) (1:2000 dilution). Protein bands were visualized by chemiluminescence using the ECL Western Blotting Substrate (Thermo Fisher Scientific, MA, USA).

### Gene regulation in cells with knocked-down RNase H1

NCI-H358 cells were seeded in 24-well plates for 12 h before transfection. The transfection medium was formulated in serum-free medium containing 5 μg/mL of Lipofectamine2k (Invitrogen) and either 200 nM of mouse small-interfering RNA (siRNA) that specifically targets the human RNaseH1 (SMARTpool, Dharmacon) or 200 nM of nontargeting control siRNA (siControl) (Dharmacon). After 4 h of transfection, the transfection medium was switched to full medium for 20 h. The cells were then cultured in RPMI-1640 medium supplemented with 0.5% FBS and 1% penicillin-streptomycin for another 48 h. Thereafter, cells were incubated with full medium and pacDNA or Lipofectamine2k-complexed PS ASOs for 6 h. Cells were harvested and digested for western blotting as indicated above. The primary antibody used was first incubated with RNase H1 polycolonal antibody (Cat. No.156061AP, Proteintech Group Inc.) (1:1000 dilution), and visualized by incubating with anti-rabbit IgG, HRP-linked secondary antibody (Cell Signalling Technology, MA, USA) (1:2000 dilution).

### Coarse-grained molecular dynamics simulation

An all-atom structure of pacDNA was mapped to coarse-grained (CG) beads according to functional groups that best match the bead types in the MARTINI force field^4^ (2-5 atoms per bead). The CG parameters for the polymer backbone and linkers were extracted from a molecular dynamics (MD) trajectory of an atomistic simulation of a three-repeating unit model molecule based on the OPLS-AA force field.^5^ The coarse-grained structure was solvated in a CG water box. Sodium ions were added to ensure the system is neutral in charge. The solvated system underwent energy minimization, followed by 50 ns of equilibration and 1 μs of production MD simulation (step size: 4 fs; NPT ensemble) using GROMACS 2021.3^6^ with the velocity rescale thermostat^7^ and the Parrinello-Rahman barostat^8^ under 300 K and 1 bar.

---

**Scheme S1.**
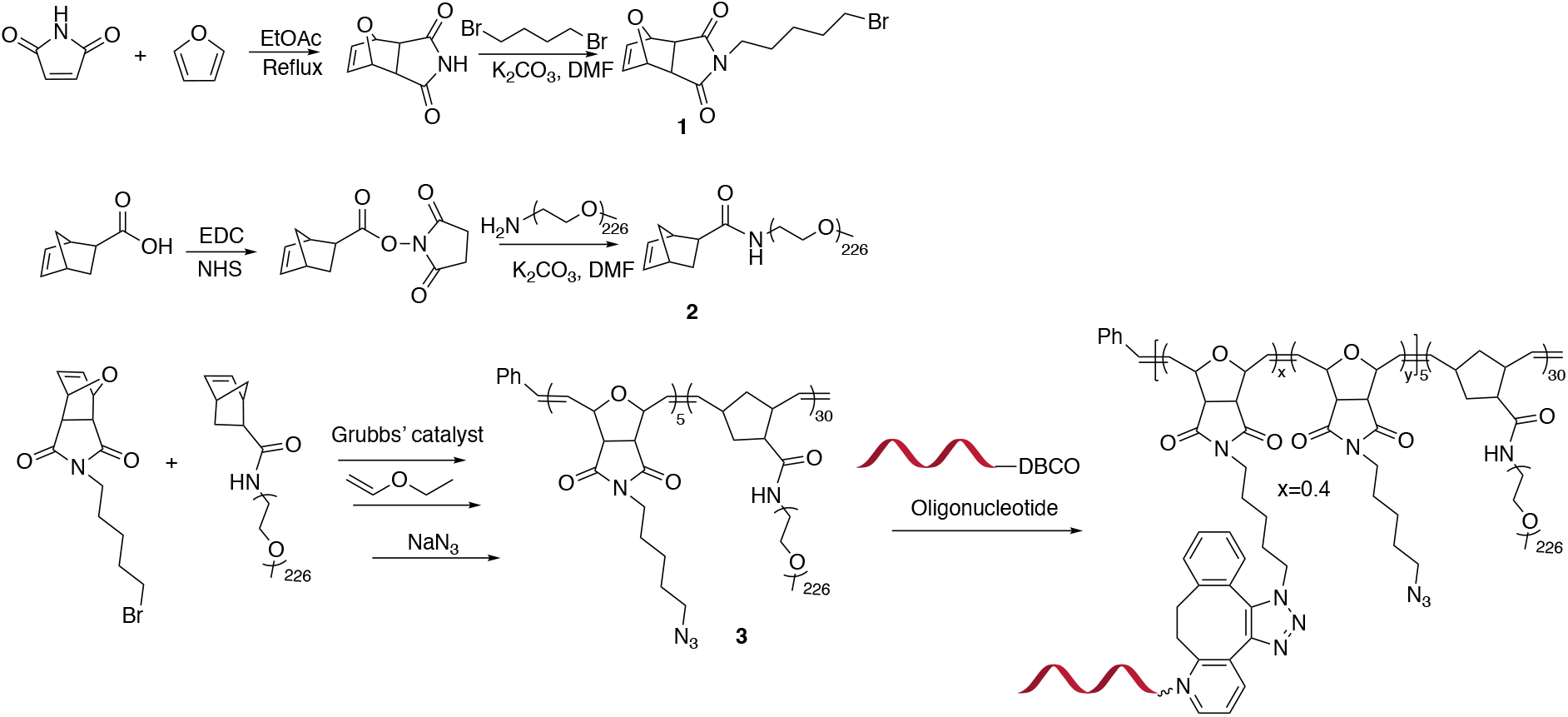
Chemical synthesis of the pacDNA.

**Table S1.**
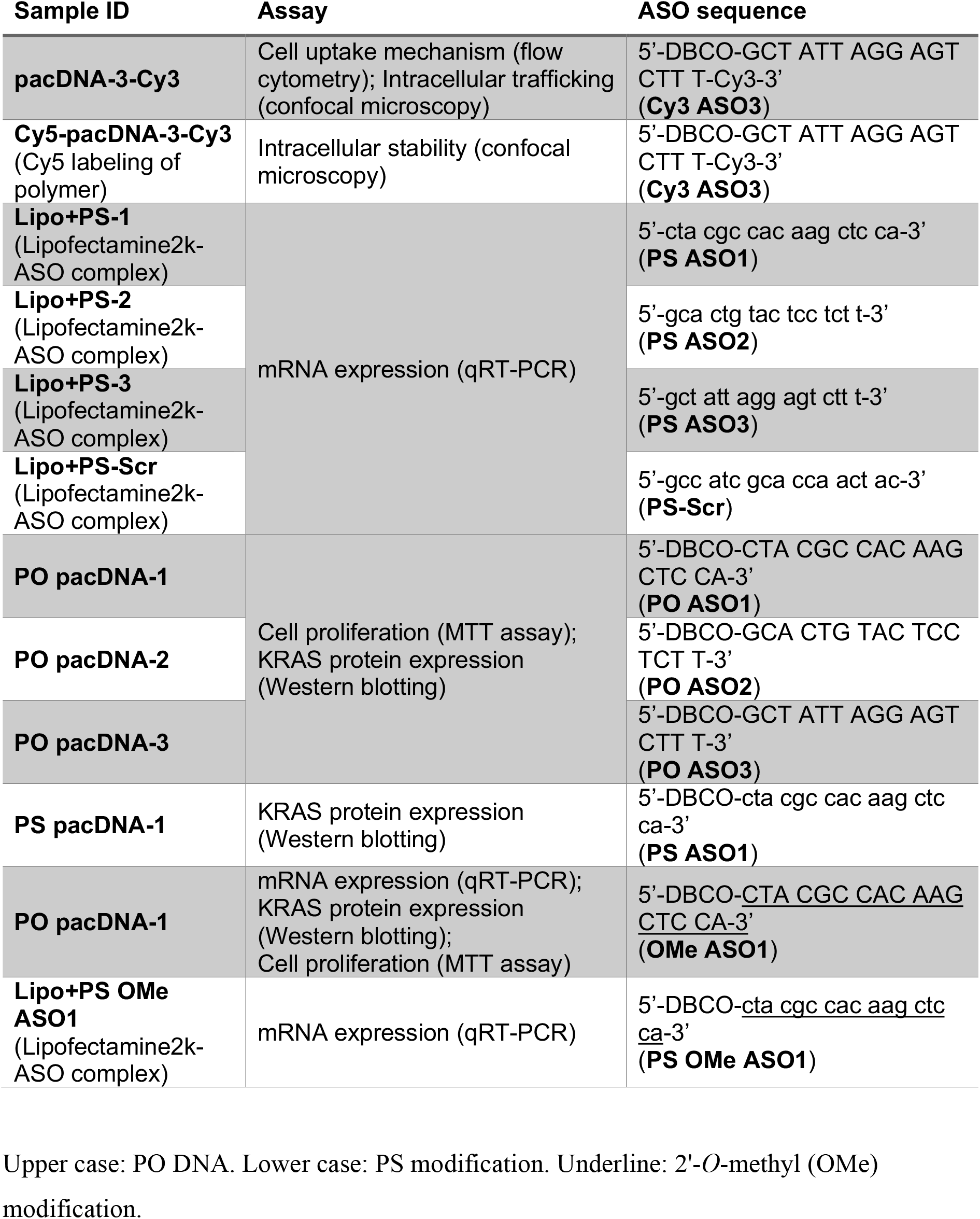
Nomenclature of samples, ASO sequence, and assay performed.

**Table S2.** List of pharmacological inhibitors of pathways of endocytosis and receptors.

| **Target** | **Agent/condition** | **Stock solution** | **Final concentration** |
| --- | --- | --- | --- |
| **Clathrin** | Chlorpromazine | 1 mg/mL (in water) | 5 µg/mL |
| **Energy-dependent endocytosis** | 4 °C | - | - |
| **Lipid raft/caveolae** | Filipin III | 2 mg/mL (in 100% DMSO), then dilute 1:7 in PBS | 2.5 µg/mL |
| **SR-A receptor** | Fucoidan | 10 mg/mL (in water) | 100 µg/mL |
| **Macropinocytosis** | Amiloride | 250 mg/mL (in 100% DMSO), then dilute 1:9 in water | 250 µg/mL |
| **Dynamin** | Dynasore | 80 mM (in 100% DMSO) | 80 µM |
| **Lipid raft/caveolae** | M-β-CD | 250 mg/mL (in water) | 12.5 mg/mL |

**Figure S1.**
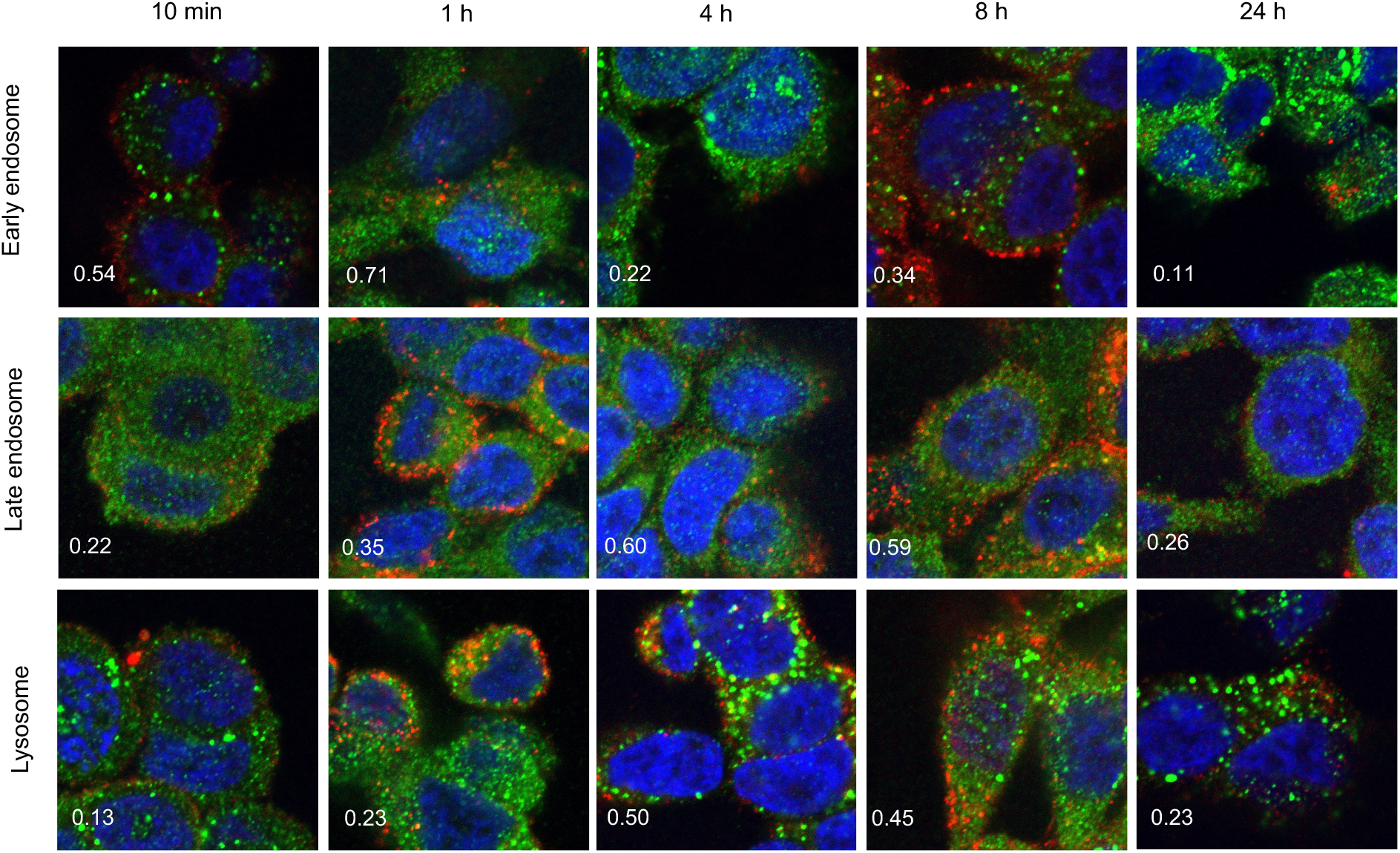
Intracellular trafficking of pacDNA in NCI-H358 cells following different durations of incubation in serum-deprived medium. pacDNA: red; immunofluorescence staining of organelle markers: green. The markers include EEA1 (early endosome), Rab9 (late endosome), and LAMP1 (lysosome). Manders’ colocalization coefficient, shown in the bottom left side of each image, of 0.5 or above indicates substantial colocalization.

**Figure S2.**
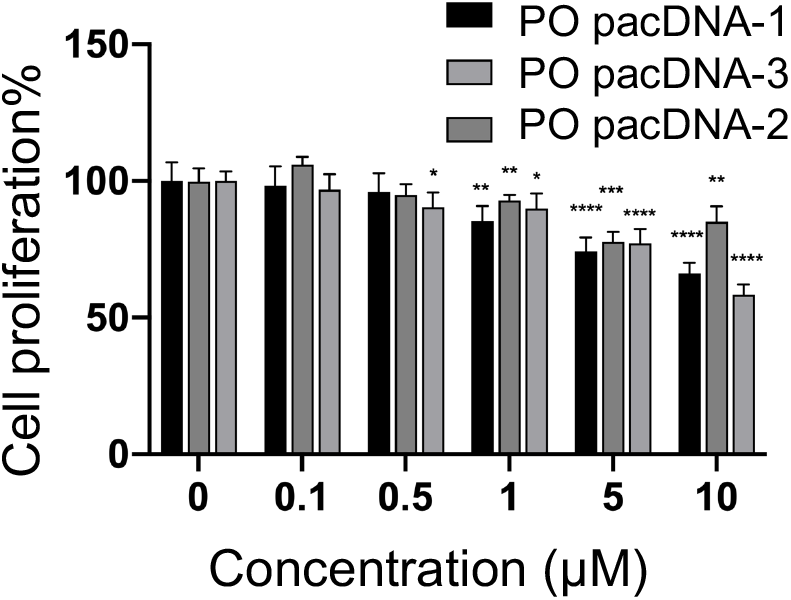
Cell viability of NCI-H358 treated with pacDNA with different antisense DNA sequences for 96 h as determined by an MTT assay. Error bars denote the standard deviation resulting from five individual experiments.

